# Policies or Knowledge: Priors differ between perceptual and sensorimotor tasks

**DOI:** 10.1101/132829

**Authors:** Claire Chambers, Hugo Fernandes, Konrad Paul Kording

## Abstract

If the brain abstractly represents probability distributions as knowledge, then the modality of a decision, e.g. movement vs perception, should not matter. If on the other hand, learned representations are policies, they may be specific to the task where learning takes place. Here, we test this by asking if a learned spatial prior generalizes from a sensorimotor estimation task to a two-alternative-forced choice (2-Afc) perceptual comparison task. A model and simulation-based analysis revealed that while participants learn the experimentally-imposed prior distribution in the sensorimotor estimation task, measured priors are consistently broader than expected in the 2-Afc task. That the prior does not fully generalize suggests that sensorimotor priors strongly resemble policies. In disagreement with standard Bayesian thought, the modality of the decision has a strong influence on the implied prior distribution.

**NEW AND NOTEWORTHY:** We do not know if the brain represents abstract and generalizable knowledge or task-specific policies that map internal states to actions. We find that learning in a sensorimotor task does not generalize strongly to a perceptual task, suggesting that humans learned policies and did not truly acquire knowledge. Priors differ across tasks, thus casting doubt on the central tenet of may Bayesian models, that the brain’s representation of the world is built on generalizable knowledge.

## INTRODUCTION

The acquisition of knowledge is thought to be at the core of the brain’s function (Tenenbaum et al. 2006, 2011; Battaglia et al. 2013). A behavioral signature of knowledge-use is strong generalization across situations. For instance, when a child learns a new word they can use it in many new situations, not just the sentence where the word was learned (Xu and Tenenbaum 2007; Perfors et al. 2011). However, the framing of learned representations as generalizable knowledge may not apply to all of the brain’s functions equally. For example, generalization from movements of one arm to those of the other is not always complete (Criscimagna-hemminger et al. 2003; Shadmehr 2004). Indeed, the reinforcement learning literature (Sutton and Barto 1998) defines an alternative way of learning. Within this framework, learning is framed as policy-acquisition, i.e. mappings from states to actions (Daw and Doya 2006; Haith and Krakauer 2013). This definition implies that learning of policies is specific to the action for which it was learned and thus suggests limited generalization across tasks. We want to know if humans are policy animals, knowledge carriers, or something in between.

In sensorimotor estimation tasks, humans weigh prior knowledge with sensory information in a near-optimal way (Körding and Wolpert 2004; Tassinari et al. 2006; Berniker et al. 2010; Vilares et al. 2012) and generalize learned prior statistics to new conditions (Fernandes et al. 2014). Thus, there is evidence for learning of sensorimotor priors. However, little is known about whether sensorimotor learning generalizes when the read-out modality of the decision changes. Therefore, we do not know if sensorimotor priors should be described as knowledge or policies. This is important because it has consequences for how neural representations should be conceptualized.

Here, we investigate if priors are the same across modalities by examining whether priors generalize across two simple tasks. The experiment was designed so that tasks were equivalent in terms of how probabilistic information should be combined to achieve optimal performance. Participants learned a spatial prior in a sensorimotor estimation task, and we asked if they transferred the learned prior to a two-alternative-forced-choice (2-Afc) task, where participants made a binary decision about object location. We inferred the standard deviation of the learned prior and found that the learned sensorimotor prior does not generalize fully to the 2-Afc task. The prior standard deviation measured from 2-Afc decisions was higher than the standard deviation measured from sensorimotor estimates. This shows that a learned prior does not generalize fully across sensorimotor and decisional modalities and suggests that sensorimotor priors are represented as policies.

## METHODS

The results presented here use data from previous work (Acuna et al. 2015), augmented with newly collected data on the same paradigm. A complete description of the methods is given in previous work and will be described here. Participants were six males and two females (age: M = 29.87, SD = 7.27). Participants gave written informed consent before taking part. Ethical approval was provided by the NU IRB #20142500001072 (Northwestern University, USA).

We required tasks that were equivalent in how probabilistic information should be combined across sources and that allowed us to infer priors used by participants. We used a “coin-catching” task (Berniker et al. 2010; Vilares et al. 2012; Acuna et al. 2015), where on each trial, participants guessed the location of a hidden stimulus (“coin”) on the screen based on an uncertain visual cue (“splash”) and a prior learned through feedback on stimulus location. Varying the prior and likelihood width allowed us to assess whether participants weighed prior and likelihood information according to their relative uncertainties during sensorimotor estimation and decision making.

Before starting the experiment, participants were presented with the instructions that on each trial, someone was throwing two coins, one after another, into a pond represented by the screen; and that their aim was to guess where the coin stimuli landed. They were told that there was no relationship between where the two coins landed (Fig. 1). On each trial, they were presented with “splash” stimuli and were told that it was caused by a hidden coin stimulus. On estimation trials, participants provided an estimate of the second stimulus’s location on the horizontal axis by placing a vertical bar where they thought that the stimulus landed. On 2-Afc trials, participants compared the locations of the inferred stimulus locations and indicated which stimulus was further to the right. Participants were paid based on their performance on the estimation task, as quantified using the distance between their estimates and the true stimulus location.

**Figure 1.**
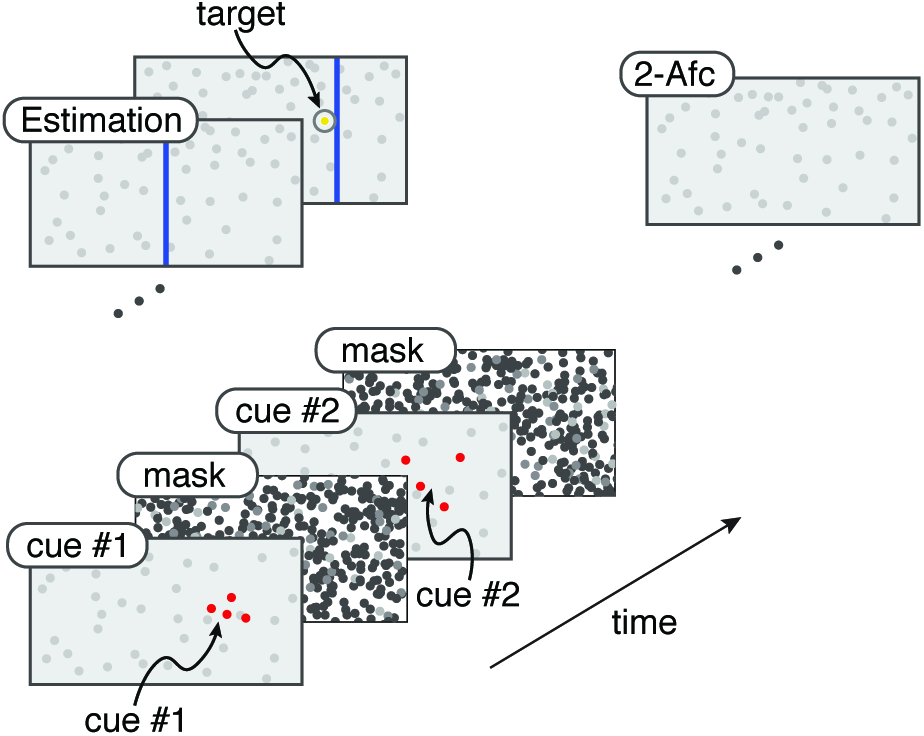
Experimental protocol. Participants were shown two splashes (likelihoods) in succession, created by hidden coins (stimuli) falling into a pond (screen), which were interleaved with white noise masks. Participants were then presented with one of two possible tasks. In the estimation task, participants were prompted to place a net where the second hidden stimulus fell. In the 2-Afc task, participants reported which hidden stimulus landed farther to the right.

Eight participants performed the experiment, including the seven participants from an existing data set (Acuna et al. 2015) and one additional participant to increase the power of group statistics (statistical results were the same with and without this participant). The experiment lasted 10,000 trials over 5 days. On each day, they were seated in front of a computer monitor (52 cm wide, 32.5 cm high) in a quiet room. Stimuli were generated by sampling visual stimuli from a Gaussian prior distribution defined over spatial location, with a mean at the center of the screen and standard deviation of .04 or .2 in units of screen width. The stimulus was hidden from view. Instead, they were presented with a visual cue with experimentally-controlled uncertainty (splash stimulus). The splash consisted of four dots sampled from a Gaussian likelihood distribution centered on the stimulus location. The likelihood distribution could have a standard deviation of .025 or .1 in units of the screen. In all trials, two consecutive splashes were displayed for .025 s, each followed by a visual mask for .5 s. The standard deviation of the likelihood was either the same across presentations within a trial (both at .025 or both at .1) or varied within trial (.025 and .1) and presented in randomized order. We refer to the broader likelihood as the reference and the narrower likelihood as the probe.

On each trial, participants performed one of two tasks, as defined by the question displayed at the end of the trial. On estimation trials, participants were asked “Where was the coin located?” and they indicated where they thought the second coin stimulus was using a vertical bar (“net”), which was 2% screen width and extended from the top to the bottom of the screen. On 2-Afc trials, participants were asked “Which coin was further to the right?” and using a key-press they indicated if they thought the first or second coin stimulus was further to the right. Trials in both tasks were identical until the end of the trial, until the question was displayed on screen. At the end of estimation trials only, feedback was provided on the exact location of the stimulus, but not on 2-Afc trials, allowing us to ask if the prior learned in the estimation task generalizes to the 2-Afc task.

### Experimental Design

There were four conditions in the estimation task: Narrow Prior, Narrow Likelihood; Narrow Prior, Wide Likelihood; Wide Prior, Narrow Likelihood; Wide Prior, Wide Likelihood. In the 2-Afc trials, conditions were defined by the width of the prior (Narrow Prior and Wide Prior) and whether likelihoods were equal within trial, Equal Likelihoods (both narrow or both wide) or Unequal Likelihoods (one narrow and one wide). We only used Unequal Likelihood trials in the present study. Therefore in our analysis, there were two conditions for the 2-Afc trials: Narrow Prior and Wide Prior.

On each day of the experiment participants performed two 1,000-trial blocks. The prior over stimulus location switched from block to block (e.g., from wide to narrow on one day, from narrow to wide on the subsequent day, and so on). Each block contained 500 estimation trials and 500 2-Afc trials in a random order all generated from the same prior. In order to aid with learning the prior, estimation trials made up the first half of each block (375 estimation trials and 125 2-Afc trials), and 2-Afc trials made up the second half of each block (125 estimation trials and 375 2-Afc trials).

### Data Analysis and simulations

We asked whether the use of prior information differed between psychophysical tasks. To answer this question, we examined whether the prior parameters fit to one of the two tasks could predict behavior well in the other task. As a baseline comparison, we examined whether the prior standard deviation fit to one half of the data predicted behavior on the other half of the data, within task. To examine how each participant’s prior related to the veridical prior used in the experiment, we estimated the prior parameters from each task. To ensure that the data analysis produced unbiased results, we performed the same analysis on data simulated from an ideal Bayesian model.

### Quantifying the Estimation slope and PSE slope from behavioral data

In order to examine the use of probabilistic information during the estimation task, we examined how much participants relied on likelihood or prior information. In the estimation task, participants gave a continuous estimate of stimulus location. We wanted to quantify how much participants relied on the learned stimulus location (prior) or visual information (likelihood). To do so, we computed the relationship between the likelihood’s center and participants’ estimates, which we termed the *Estimation slope.* If someone were to rely only on likelihood information to judge stimulus location, on average, estimates should correspond to the center of the likelihood (*Estimation slope* =1). If someone were to ignore the likelihood entirely and rely only on their prior to judge stimulus location, there should be no relationship between the likelihood’s center and estimates (*Estimation slope* = 0). The Bayesian optimal strategy is to weigh the prior and likelihood according to their relative precision, as in Equation 1.

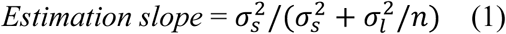

In the 2-Afc task, participants were given probabilistic information on stimulus location exactly as in the estimation task. On each trial, they compared the locations of two stimuli with different uncertainties, a probe stimulus with Narrow Likelihood and a reference stimulus with Wide Likelihood. Uncertainty should influence the judgment of stimulus location in the same way as in the estimation task. A Bayesian observer judges the more uncertain stimulus to be shifted further to the prior mean than the more certain stimulus. This, in turn, influences decisions about relative stimulus location. Therefore, use of the prior can be inferred from participants’ 2-Afc data.

Consider the psychometric function that describes the comparison of stimuli with unequal widths. The psychometric function is the probability that the probe stimulus is reported to the right, *P(Decision=1)*, as a function of difference between the likelihood stimuli (*Discrepancy*), and the *Reference location*. The participant’s prior influences the *Discrepancy* at which the stimuli are perceived as equal (point of subjective equality, *PSE*). For a Bayesian observer, the *PSE* arises when the reference is more distant from the prior’s center than the probe. The *PSE* further deviates from zero as the distance between the prior and the reference increases. Importantly, the slope of this linear relationship, the *PSE slope*, is related to the width of the participant’s prior – a *PSE slope* of 0 shows that participants relied only on visual information from the likelihood; and the more negative the *PSE slope*, the narrower the participant’s prior. The optimal *PSE slope* is given by Equation 2.

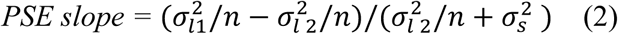

We fit psychometric functions (the cumulative Gaussian function) to each participant’s decision data. The *PSE slope* (*m*_*PSE*_) estimated from this function provides an indicator of the variance of the participant’s prior. We model the probability of a decision as:

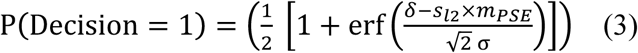

where *δ* is the discrepancy between stimuli, *s*_*l*2_ is the location of the reference stimulus with broader likelihood, *σ* describes the deviation of the function (Acuna et al. 2015). We find the values of *m*_*PSE*_ and *σ* using a maximum-likelihood estimation algorithm.

### Analysis of priors during Estimation and 2-Afc decision making

If priors are the same, then priors used in one task should predict behavior in the other task well. A cross-validation error computed across tasks should not exceed the error computed within tasks. To test this, we performed 2-fold cross-validation by estimating priors from one task and computing the Mean Squared Error (MSE) on the held-out task (Across-task MSE). We compared the Across-task MSE with the within-task MSE, computed by performing 2-fold cross-validation using the data of one task, then summing the MSE across tasks. If priors are the same, we expect that the Across-task MSE should not exceed the Within-task MSE. This analysis allowed us to examine if priors were the same or different across tasks.

To quantify the prior width in the estimation task and the 2-Afc task, we used the *Estimation slope* and the *PSE slope* respectively. Using a maximum-likelihood estimation algorithm, we estimated prior standard deviation parameters from the slopes of one task by minimizing the MSE between the slope values and the slopes given by Equations 1 and 2. To compute the Across-task MSE, we predicted the slopes of the held-out task from the fitted parameters and compared the predicted slope with that computed from the data using the MSE. To compute the Within-task MSE, we estimated the prior standard deviation parameters from 50% data of one task, then predicted the slope for the same task, and compared the predicted slope to the slope computed from the held-out data.

To ensure that our analysis led to unbiased results, we simulated 1000 Bayesian observers who combined prior and likelihood information optimally and used the veridical prior parameters in both tasks. Simulated observers should not show systematic differences between the Across-task and Within-task MSE.

In order to examine how the priors used by participants differed from the veridical priors, we estimated the prior standard deviation using Equations 1 and 2 and a maximum likelihood estimation algorithm. We ensured that this procedure did not lead to biased results using simulations. We simulated Bayesian observers who combined prior and likelihood information optimally. Simulated participants used the veridical prior standard deviation in the estimation task and used a prior standard deviation in the 2-Afc task which related to the veridical value by a factor of .5, 1, or 2. We performed 1000 simulations per condition. Inferring the prior width from the behavioral data allowed us to examine generalization of the prior.

## RESULTS

We asked if a learned prior distribution generalizes across tasks and thus consists of knowledge. To do so, we first had participants learn a prior in a sensorimotor estimation task where participants gave a continuous estimate of stimulus location under uncertainty. We then quantified use of the prior in a 2-Afc task where instead participants compared the locations of two hidden stimuli (Fig. 1). We used data from previous work (Acuna et al. 2015). We examined whether the data was consistent with use of the same or different priors across tasks and estimated the prior standard deviation parameters from the data of each task.

In our tasks, participants judged the location of visual stimuli on screen. Stimuli were samples from a Gaussian prior distribution, 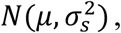, which were hidden from view. Instead, participants were shown an uncertain visual cue in the form of *n* samples (*n*=4) from a Gaussian likelihood distribution, distributed around stimulus location, 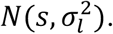 When judging stimulus location, the Bayesian optimal strategy is to combine the likelihood and the learned prior according to their relative precision. Therefore, to examine participants’ use of probabilistic information, we manipulated the standard deviations of the prior and likelihood, 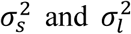 and quantified how much participants rely on the likelihood or prior to reach a decision (see Methods, Fig. 2). Our paradigm allowed us to examine integration of probabilistic information and to infer participants’ learned priors in the estimation and 2-Afc tasks.

**Figure 2.**
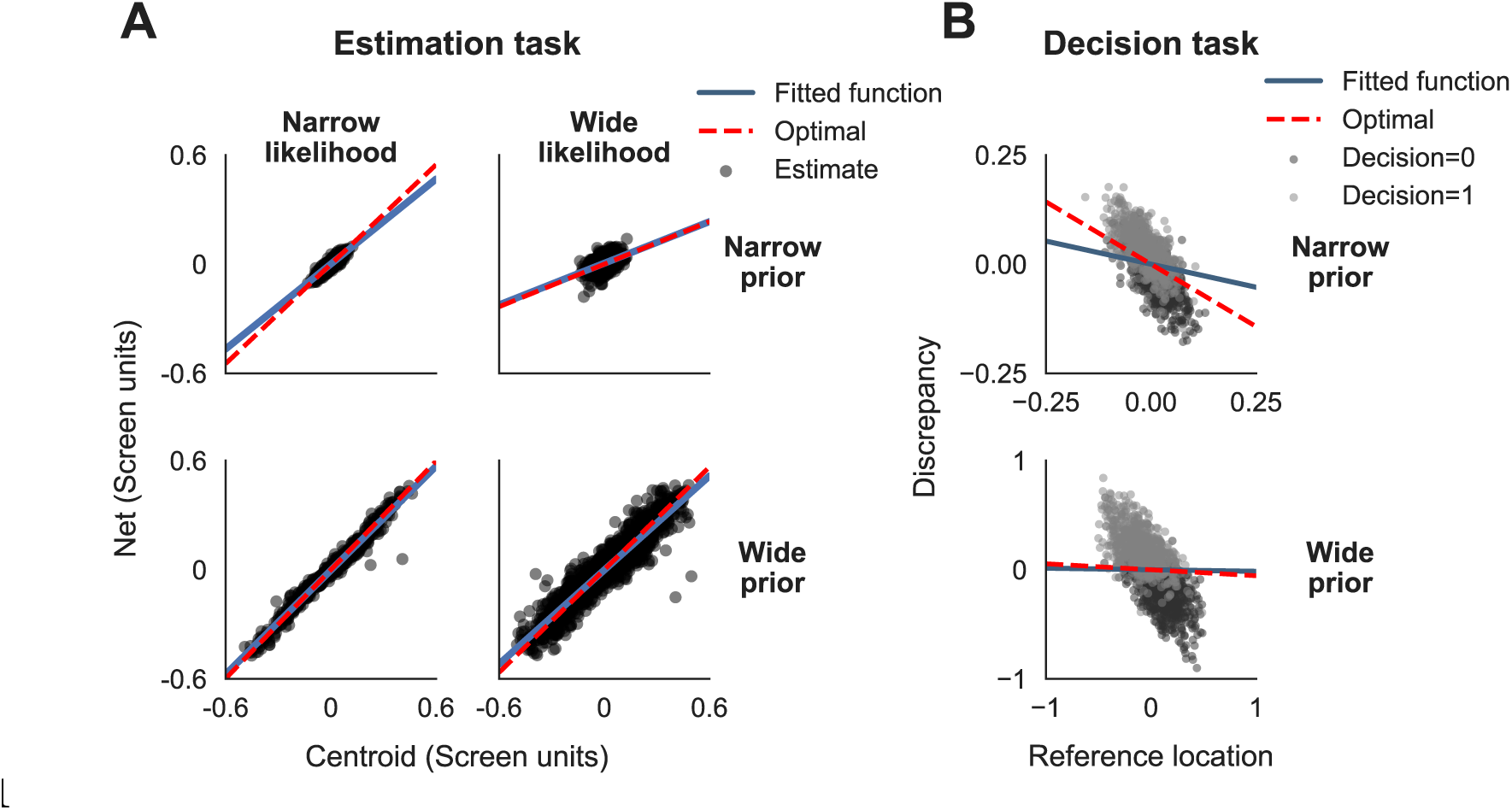
Estimation and 2-Afc data. (**A**) Estimation data overlaid with linear fit for a representative participant. The net position as a function of the centroid of the likelihood is shown for each trial (black points). Each panel displays estimation data for one condition, with overlaid fitted (blue line) and optimal (red line) functions. An *Estimation slope* of 1 indicates complete reliance on the likelihood and an *Estimation slope* of 0 indicates complete reliance on the prior. (**B**) 2-Afc data for the representative participant in (**A**) with one panel per condition. Raw binary decision data (dark gray points, Decision=0, probe stimulus to the left; light gray points, Decision=1, probe stimulus to the right). The best fitting PSE (blue line) and optimal PSE (red line) are shown. The more negative the *PSE slope*, the narrower the prior.

To examine use of probabilistic information in producing estimates, we first examined influence of the prior width and likelihood width on participants’ reliance on the likelihood or prior (*Estimation slope*, see Fig. 2 and Methods for details). We found that both prior width and likelihood width influence the *Estimation slope* (Repeated-measures ANOVA: main effect of prior width: p<.0001, F(1, 7) = 320.74; main effect of likelihood width, p<.0001, F(1, 7) = 140.67). Therefore, participants use the prior and likelihood widths to judge stimulus location. Therefore, it makes sense to describe participants’ sensorimotor estimates as Bayesian and to quantify the prior used during the task.

We then examined use of probabilistic information in the 2-Afc task, with a measure of reliance on prior or likelihood in decision data, which we termed the *PSE slope* (see Fig. 2 and Methods for details). It was important to establish that participants could incorporate a prior into 2-Afc decisions, as shown by a negative *PSE slope*. We thus compared PSE slopes with 0 (Narrow prior: p < .05, one-sample, 2-sided t-test, t(7) = 3.60, Wide prior: p < .05, one-sample, 2-sided t-test, t(7) = 3.28, Bonferroni-corrected p-values). As is shown by the significantly negative PSE slope in both conditions, participants incorporate priors into 2-Afc decisions. There was a significant effect of prior width on *PSE slope* (p < .0001, paired, 2-sided t-test, t(14) = 7.59). Thus, participants are influenced by the prior in their decisions and have a greater reliance on the prior in the Narrow-Prior condition. This is consistent with Bayes task.

**Figure 3.**
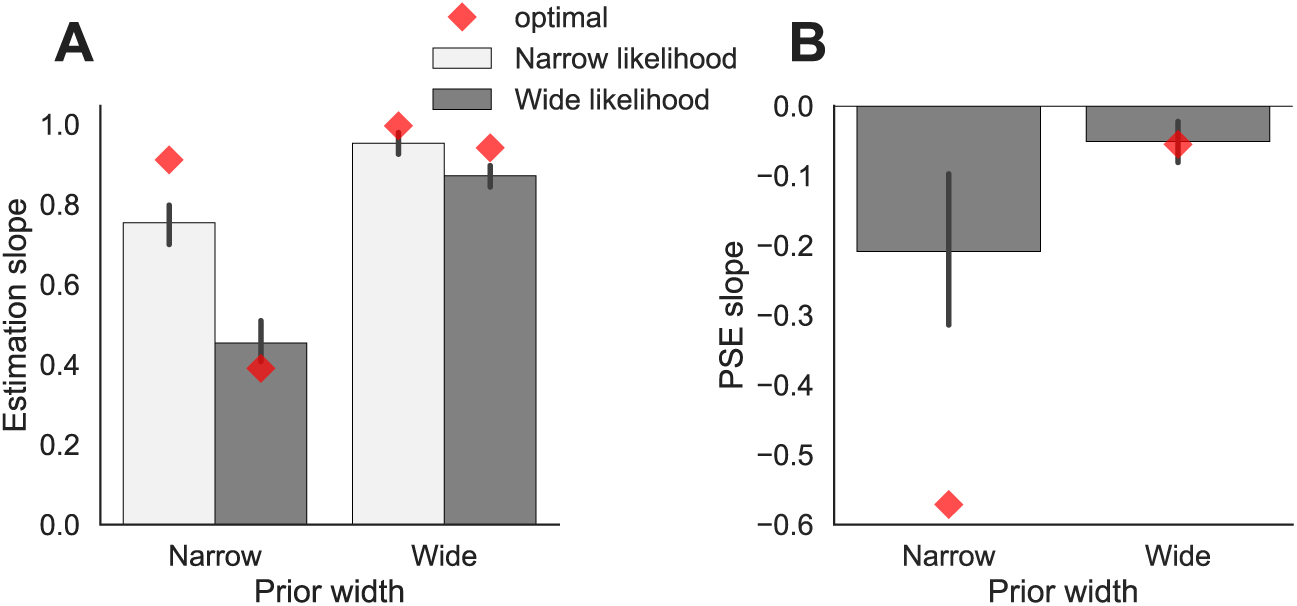
*Estimation slope* and *PSE slope*. (**A**) The median *Estimation slop*e is shown as a function of Prior width and Likelihood width. Error bars display bootstrapped 95% confidence intervals (CI). The optimal slope values for each condition are shown by red diamonds. (**B**) The median *PSE slope* is shown as a function of Prior width. Error bars display 95% CI. The optimal slope values for each condition are shown by red diamonds.

Behavior in the two tasks is in accordance with the use of probabilistic information. This was shown by an influence of the uncertainty of the prior and likelihood on judgments in both tasks. However, it is possible that the priors used in the estimation and 2-Afc tasks are different. Such a difference would be in violation of standard Bayesian thought where the prior representation is considered as knowledge and hence, domain general and fully available for use across tasks.

We then asked if the data supports task-dependent prior representations. If participants use the same prior to perform both tasks, prior width parameters estimated from one task’s data should predict the other task’s data well. The cross-validated error between slopes across tasks (Across-task MSE, see Methods) should not exceed the cross-validated error within tasks (Within-task MSE). We found that the Across-task MSE exceeded the MSE computed within each task (p<.01, t(7)=3.35, Fig 4A). Simulations show that this analysis is unbiased and does not favor this result (Fig 4A). This result suggests that participants use different priors in the different tasks.

**Figure 4.**
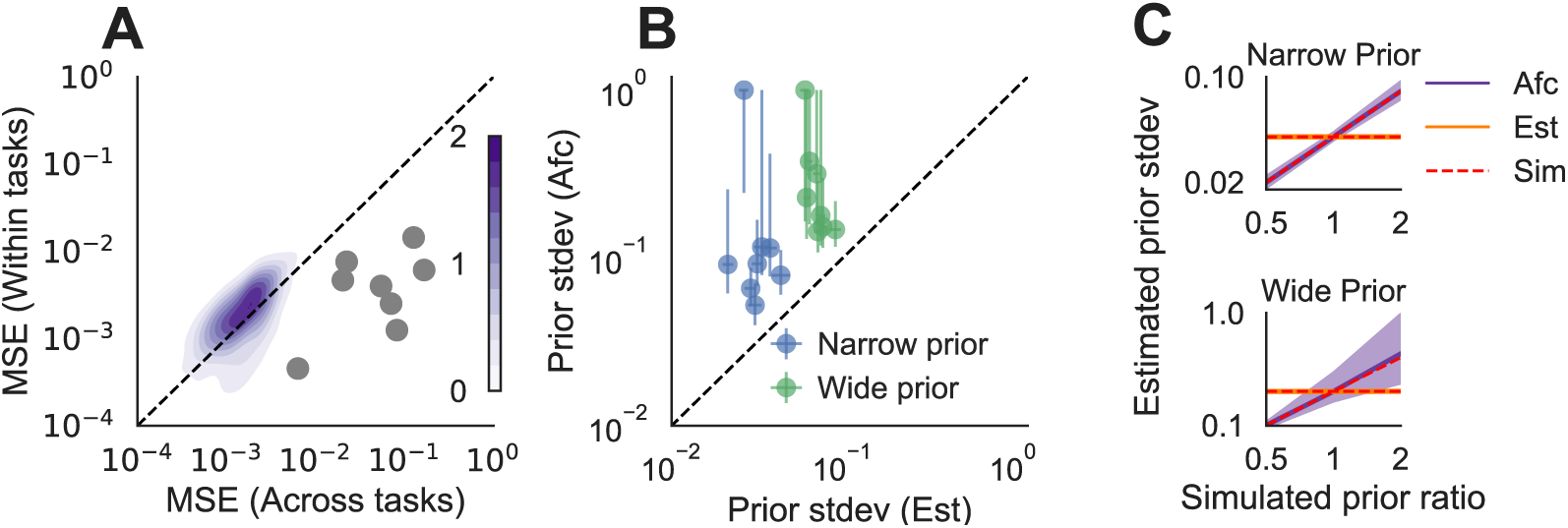
Comparison of priors in the Estimation and 2-Afc tasks (**A**) The MSE computed within tasks is shown as a function of the MSE computed across tasks. For each participant, the MSE across tasks exceeds the MSE within tasks. Therefore, the data is not consistent with use of the same prior. MSE for 1000 simulated participants (distribution in purple, color bar displays kernel density estimate) show that this analysis gives unbiased results. (**B**) For each participant, prior standard deviation inferred from the 2-Afc task data is shown as a function of the prior standard deviation inferred from the estimation task data. The median bootstrap is shown (error bar=95% CI). The dotted line shows the diagonal, for which prior width in the tasks are equal. (**C**) Prior parameters estimated from the data of 1000 simulated Bayesian observers in the Narrow-Prior condition (upper panel) and Wide-Prior condition (lower panel). Simulated participants use the theoretical prior in the estimation task and either the same prior standard deviation in the 2-Afc task (simulated prior ratio=1, the prior ratio being the ratio of the standard deviations in the 2-Afc and estimation tasks) or a different prior standard deviation in the 2-Afc task (simulated prior ratio= .5, or 2. The median inferred prior standard deviation is shown for the estimation task (orange) and the 2-Afc tasks (purple), shaded area= 2.5^th^-97.5^th^ percentile. Broken red lines show the veridical prior standard deviations.

Having found that priors were different across tasks, we wanted to know how they were different. We, therefore, inferred the prior width (standard deviation) from the estimation and 2-Afc data from the *Estimation slope* and the *PSE slope* using bootstrapped parameter estimation. For each individual participant, the prior in the 2-Afc task is wider than the prior in the estimation task in both the Narrow and Wide prior conditions (95% CI consistently above the diagonal in Fig. 4B). We show that our estimation of prior width is unbiased and that we can successfully infer prior width from simulated estimation and 2-Afc data (Fig. 4C). Regarding our hypothesis on the generalization of the learned prior from the estimation task to the 2-Afc task, this shows that the prior does not generalize fully. Therefore, our analysis supports policy representation rather than knowledge representation.

## DISCUSSION

We examined if a prior distribution learned during a sensorimotor estimation task generalized to a computationally-equivalent 2-Afc decision task. We showed that there was a difference in priors across tasks. The finding of a wider prior in the 2-Afc task shows that the prior did not generalize fully from the situation where participants provided a continuous estimate of location to different task where participants compared two object locations. This shows that sensorimotor priors are not knowledge, in the sense that they do not generalize fully across modalities.

A caveat is that we assume that the brain uses maximum a-posteriori (MAP) to compute decisions. MAP is widely-used in the decision-making literature and is a plausible choice of mechanism since it maximizes reward in simple cases (Maloney 2002; Mamassian et al. 2002). Other decision-making mechanisms include sampling from probability distributions and have been explored in previous work (Vul et al. 2014; Acuna et al. 2015). While the choice of MAP may be reasonable in the case of unimodal Gaussian posterior distributions as in the current study, MAP is less adapted to cases of multimodal or broadly-distributed posteriors. Further work is needed to explore the decision rules that the brain uses.

One implication of our finding is that priors cannot be assumed to generalize even when the difference between learning and testing conditions or tasks is subtle. For example, previous work investigating decision-making mechanisms quantifies the prior in an estimation task and measures the influence of the subjective prior in a 2-Afc task (Acuna et al. 2015). The findings of this previous work therefore rest on the assumption that the prior is the same across tasks and the conclusions of this paper and others with the same assumption should be revisited.

Why are the priors different? The tasks may engage distinct neural systems, with the estimation task having a stronger sensorimotor component (‘Where is the object in relation to me?’), whereas the 2-Afc task is a perceptual task and concerns relationships between objects in the outside world (‘Where is one object in relation to another?’). Therefore, partly independent neural representations may lead to incomplete generalization across tasks (Aglioti et al. 1995; Knill 2005). In this view, partial generalization comes from partly distinct neural systems.

Importantly, our finding is inconsistent with the view that the brain acquires fully generalizable knowledge, in the form of priors that can automatically be incorporated into behavior regardless of the task. While high-level conceptual representations may fit the definition of knowledge (Perfors et al. 2011; Tenenbaum et al. 2011; Battaglia et al. 2013), our findings show that learning in a sensorimotor task has a strong policy component, with a prior being partly confined to the task where it was learned. In naturalistic situations, the use of policies may be functionally beneficial, allowing for learning to be optimized for the task at hand.

Knowledge and policies are often evoked to explain behavior (Tenenbaum et al. 2011; Haith and Krakauer 2013). However, they are seldom pitted against each other as they originate from distinct theoretical frameworks. A more common dichotomy is that of procedural and declarative knowledge, which describes knowledge of how to perform some action and knowledge of concepts, respectively, or ‘knowing how and knowing that’ (Ryle 1945; Winograd 1975; Squire 2004). While these resemble the concepts of knowledge and policy, the declarative-procedural dichotomy does not have the same implications for generalization. Declarative knowledge is by definition generalizable, while procedural knowledge can generalize strongly or not, that is, can be consistent with knowledge or policy. Therefore, these dichotomies do not completely overlap with one another. A second common dichotomy is that of model-based and model-free behavior (Sutton and Barto 1998; Daw and Doya 2006; Doll et al. 2012). Model-based behavior leverages a model of a situation to attain a goal, while model-free behavior involves repetition of previously successful actions. When applied to our paradigm, one could conclude that the more optimal prior use in the estimation task is based on a better model of how stimuli were generated and that deviations from this in the 2-Afc task imply weaker use of a model. Our findings, however, do not support pure model-based or model-free behavior in either task and our experiment and findings are more amenable to a probabilistic treatment and quantification of priors. Discussion of findings in light of different approaches and frameworks is helpful and will be necessary to build a more unified theory of the brain’s function.

These results are compatible with a learning framework, rather than a high-level Bayesian view of the brain’s computations, where one set of priors (knowledge) is used for different output behaviors. Multi-layer neural networks provide a flexible way of modeling diverse kinds of behavior based on function optimization (LeCun et al. 2015; Marblestone et al. 2016). Within a broader network, sub-networks that implement specialized learning could produce patterns of generalization or non-generalization across conditions and tasks. Importantly, a system that learns by gradient descent will approximate Bayesian behavior without explicitly implementing Bayesian computations (Weisswange et al. 2011; Mandt et al. 2017), simply because it is the optimal strategy for estimation under uncertainty. Our finding thus casts doubt on the view that Bayesian computation is at the core of the neural code (Zemel et al. 1998; Ma et al. 2008).

## GRANTS

This work was funded by grant by NIH grant 5R01NS063399-08, awarded to KPK.

## REFERENCES

Acuna DE, Berniker M, Fernandes HL, Kording KP. Using psychophysics to ask if the brain samples or maximizes. J Vis 15: 7, 2015.

Aglioti S, DeSouza JFX, Goodale MA. Size-contrast illusions deceive the eye but not the hand. Curr Biol 5: 679–685, 1995.

Battaglia PW, Hamrick JB, Tenenbaum JB. Simulation as an engine of physical scene understanding. Proc Natl Acad Sci U S A 110: 18327–32, 2013.

Berniker M, Voss M, Kording K. Learning priors for bayesian computations in the nervous system. PLoS One 5: 1–9, 2010.

Criscimagna-hemminger SE, Donchin O, Gazzaniga MS, Shadmehr R, Sarah E, Donchin O, Michael S. Learned Dynamics of Reaching Movements Generalize From Dominant to Nondominant Arm. J Neurophysiol 89: 168–176, 2003.

Daw ND, Doya K. The computational neurobiology of learning and reward. Curr Opin Neurobiol 16: 199–204, 2006.

Doll BB, Simon DA, Daw ND. The ubiquity of model-based reinforcement learning. Curr Opin Neurobiol 22: 1075–1081, 2012.

Fernandes HL, Stevenson IH, Vilares I, Kording KP. The generalization of prior uncertainty during reaching. J Neurosci 34: 11470–84, 2014.

Haith AM, Krakauer JW. Theoretical models of motor control and motor learning. In: Routledge handbook of motor control and motor learning. London: Routledge, 2013, p. 7–28.

Knill D. Reaching for visual cues to depth: the brain combines depth cues differently for motor control and perception. J Vis 5: 103–115, 2005.

Körding KP, Wolpert DM. Bayesian integration in sensorimotor learning. Nature 427: 244–247, 2004.

LeCun Y, Yoshua B, Geoffrey H. Deep learning. Nature 521: 436–444, 2015.

Ma WJ, Beck JM, Pouget A. Spiking networks for Bayesian inference and choice. Curr Opin Neurobiol 18: 217–222, 2008.

Maloney LT. Statistical decision theory and biological vision. In: Perception and the physical world, edited by D H, R M. New York: Wiley, 2002, p. 145–189.

Mamassian P, Landy M, Maloney LT. Bayesian Modelling of Visual Perception. In: Probabilistic models of the brain: Perception and neural function, edited by Rao N, Olhausen B, Lewicki M. Cambridge, MA: MIT Press, 2002, p. 13–36.

Mandt S, Hoffman MD, Blei DM. Stochastic Gradient Descent as Approximate Bayesian Inference. arXiv: 1–30, 2017.

Marblestone AH, Wayne G, Kording KP. Toward an Integration of Deep Learning and Neuroscience. Front Comput Neurosci 10: 94, 2016.

Perfors A, Tenenbaum JB, Regier T. The learnability of abstract syntactic principles. Cognition 118: 306–338, 2011.

Ryle G. Knowing How and Knowing That?: The Presidential Address. Proc Aristot Soc 46: 1–16, 1945.

Shadmehr R. Generalization as a behavioral window to the neural mechanisms of learning internal models. Hum Mov Sci 23: 543–568, 2004.

Squire LR. Memory systems of the brain: A brief history and current perspective. Neurobiol Learn Mem 82: 171–177, 2004.

Sutton RS, Barto AG. Reinforcement learning: An introduction. Cambridge, MA: MIT press, 1998.

Tassinari H, Hudson TE, Landy MS. Combining priors and noisy visual cues in a rapid pointing task. J Neurosci 26: 10154–63, 2006.

Tenenbaum JB, Griffiths TL, Kemp C. Theory-based Bayesian models of inductive learning and reasoning. Trends Cogn Sci 10: 309–318, 2006.

Tenenbaum JB, Kemp C, Griffiths TL, Goodman ND. How to grow a mind: statistics, structure, and abstraction. Science 331: 1279–1285, 2011.

Vilares I, Howard JD, Fernandes HL, Gottfried JA, Kording KP. Differential representations of prior and likelihood uncertainty in the human brain. Curr Biol 22: 1641–1648, 2012.

Vul E, Goodman N, Griffiths TL, Tenenbaum JB. One and done? Optimal decisions from very few samples. Cogn Sci 38: 599–637, 2014.

Weisswange TH, Rothkopf CA, Rodemann T, Triesch J. Bayesian Cue Integration as a Developmental Outcome of Reward Mediated Learning. 6, 2011.

Winograd T. Frame representations and the declarative/procedural controversy. In: Representation and understanding: Studies in cognitive science, edited by Bobrow J. Elsevier, p. 185–210.

Xu F, Tenenbaum JB. Word Learning as Bayesian Inference. Psychol Rev 114: 245–250, 2007.

Zemel RS, Dayan P, Pouget A. Probabilistic interpretation of population codes. Neural Comput 10: 403–430, 1998.

